# A spatial continuum of tumor cell states links invasion to lymphovascular dissemination in inflammatory breast cancer

**DOI:** 10.64898/2026.11.12.765468

**Authors:** David P. Mysona, Pritha Pai, Wendy A. Woodward, Jadee L. Neff, Stephen D. Gottschalk, Ashley Fletcher, Ifeoluwatomi O. Fagbemi, Gayathri R. Devi, The MDACC Inflammatory Breast Cancer (MDACC-IBC) Registry and DI Biopsy Team

## Abstract

Inflammatory breast cancer (IBC) is an aggressive breast cancer subtype marked by extensive lymphovascular invasion (LVI) of tumor cell clusters termed tumor emboli and early metastatic dissemination. The molecular adaptations that enable IBC tumor cells to survive across distinct microenvironmental niches remain poorly understood. Here, whole-transcriptome spatial profiling of human IBC specimens defines epithelial tumor cell states in the tumor stroma, leading edge, and LVI compartments relative to adjacent normal tissue. LVI tumor emboli exhibit altered programs involving antigen processing and presentation, apoptotic signaling, cell adhesion, and cholesterol biosynthesis. Using a tumor-lymphatic interface biomimetic platform (T-LAB), we show that disseminating tumor cells recapitulate key LVI-associated programs and conserved adaptive stress response (ASR) genes involving oxidative stress, NF-κB, JAG1-Notch, and XIAP signaling. Furthermore, T-LAB-derived circulating cells retain a pharmacologically actionable vulnerability to oxidative stress, linking conserved adaptive stress response remodeling to a potential therapeutic opportunity during lymphatic dissemination.

**Highlights:**

1. IBC tumor emboli exhibit distinct spatially defined molecular programs across the tumor stroma, leading edge, and lymphovascular invasion (LVI) compartments.
2. LVI tumor emboli undergo molecular adaptations involving antigen processing and presentation, apoptotic signaling, cell adhesion, and cholesterol biosynthesis.
3. The tumor-lymphatic interface biomimetic platform (T-LAB) recapitulates key LVI-associated transcriptional programs observed in human IBC specimens.
4. Lymphatic dissemination activates conserved adaptive stress response (ASR) pathways, including oxidative stress, NF-κB, JAG1-Notch, and XIAP signaling.
5. Disseminating tumor cells retain an actionable vulnerability to oxidative stress, revealing a potential therapeutic opportunity to target IBC during lymphatic dissemination.

**Graphical abstract:** 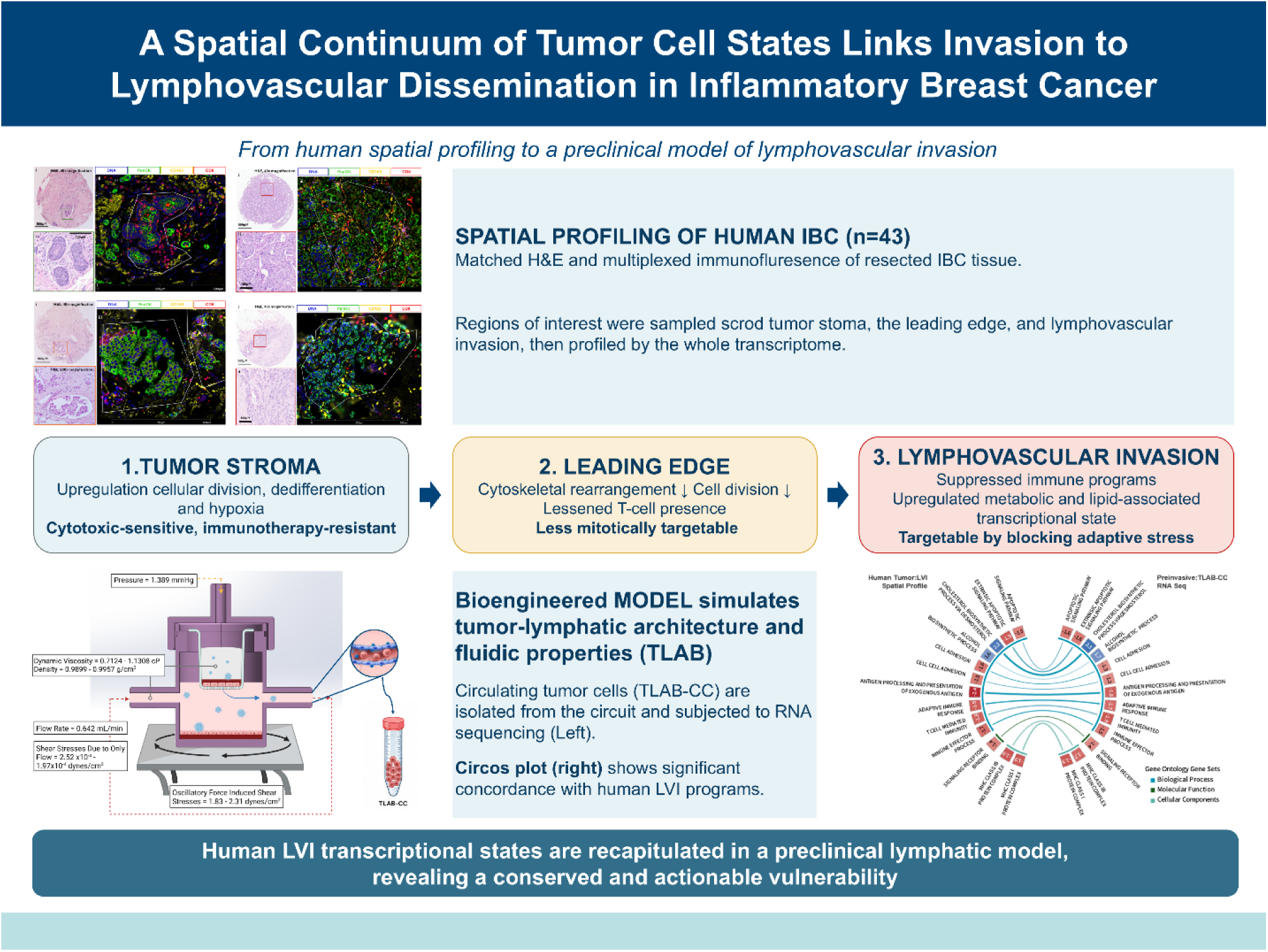

## Introduction

Inflammatory breast cancer (IBC) is a rare but highly aggressive subtype of breast cancer characterized by rapid onset, early metastatic dissemination, and disproportionately poor survival compared with non-inflammatory breast cancers^1,2^. Clinically, IBC presents with diffuse erythema or discoloration, edema, and peau d’orange of the breast, features driven in part by extensive tumor emboli within dermal lymphovascular spaces^3,4^. Despite multimodality treatment, including systemic therapy, surgery, and radiation, long-term outcomes remain poor, with 5-year overall survival rates substantially lower than those observed in other locally advanced breast cancers. Limited awareness of this rare subtype^5–7^, coupled with its aggressive clinical course, underscores the need to elucidate the biological mechanisms driving early dissemination and therapeutic resistance.

Local tumor invasion and lymphovascular invasion (LVI) within the breast and overlying skin are critical steps in the IBC metastatic cascade and, as in other malignancies, are strongly associated with poor clinical outcomes^8,9,10,11,12^. Despite its clinical significance, the molecular adaptations that support tumor cell survival and dissemination within lymphovascular spaces remain poorly understood, in part because these tumor emboli are difficult to isolate and study directly in human tumors^13,14^.

IBC represents a particularly compelling context in which to study lymphovascular biology because lymphatic tumor emboli are defining histopathologic hallmarks of the disease^2,15–17^. Unlike many models of metastasis that emphasize single-cell dissemination through epithelial-to-mesenchymal transition, IBC is characterized by cohesive clusters of E-cadherin–positive tumor cells believed to invade collectively through lymphovascular channels^14,18,19^. Prior work has identified signaling pathways associated with IBC aggressiveness^1,20–22^, including inflammatory cytokine signaling^23–26^, cholesterol metabolism^27^, and cellular stress responses^28–31^, but the spatially resolved transcriptional programs that distinguish tumor cells in lymphovascular spaces from those in the surrounding tumor stroma remain unknown.

In the present study, we used whole-transcriptome digital spatial profiling of residual human IBC post-chemotherapy specimens to compare epithelial tumor cells in stromal and lymphovascular compartments. We hypothesized that tumor cells within lymphovascular spaces adopt distinct transcriptional programs that facilitate immune evasion and metastatic dissemination. Our findings revealed that lymphovascular tumor emboli that are not eradicated by neoadjuvant systemic chemotherapy exhibit unique suppression of antigen presentation while the leading edge of the tumor produces TGFβ and undergoes cytoskeletal rearrangement. Subsequently, using a tumor–lymphatic interface biomimetic platform (T-LAB)^32^, we show that disseminating tumor cells recapitulate key LVI-associated programs and conserved adaptive stress response (ASR) identified in human IBC LVI. Importantly, this LVI-associated state retained a therapeutically actionable vulnerability, establishing T-LAB as a functional model to interrogate tumor-cell adaptation and therapeutic strategies to prevent lymphatic dissemination.

## Results

### Spatial Context Identifies Conserved Transcriptomics in Tumor Compartments

To define epithelial transcriptional programs associated with tumor stroma, leading edge, and lymphovascular compartments in IBC, we performed whole-transcriptome Digital Spatial Profiling on epithelial regions from tumor stroma, leading edge, LVI, and adjacent normal tissue across 43 IBC specimens (**Table 1**; **Figure 1A**). Regions of interest were selected on H&E-stained tissue microarrays by a pathologist, and epithelial cells were segmented using PanCK immunofluorescence (**Figure 1**). Examples from one IBC patient illustrate the study design (**Figure 1B-E**).

**Figure 1.**
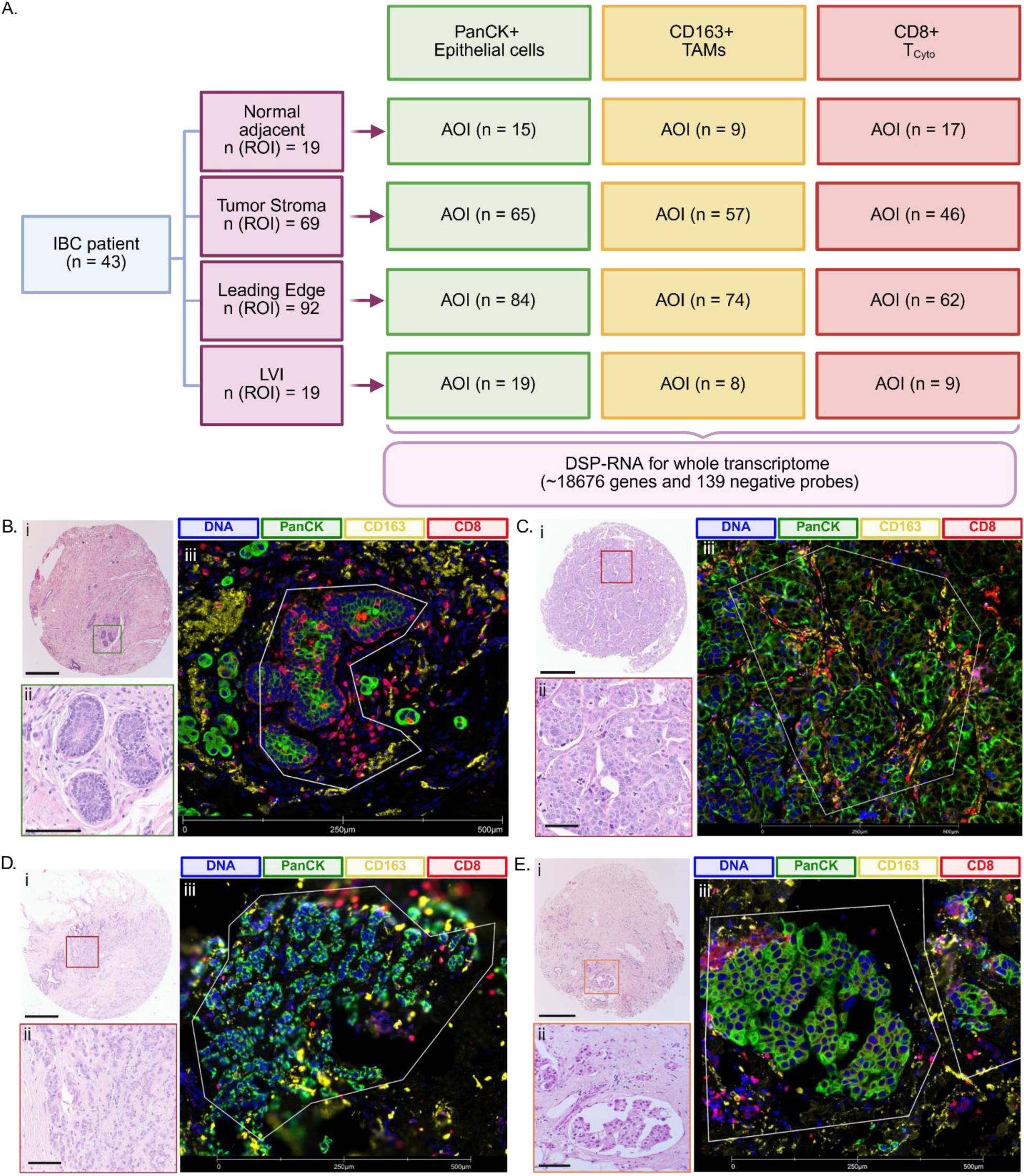
DSP-transcriptomic profiling of IBC patient samples. **A.** Schematic representation of ROIs and AOIs for whole transcriptome analysis as collected. Normal adjacent (**B**), Tumor – middle (**C**), Tumor – Edge (**D**) and LVI (**E**) ROI at 40X magnification (i, Scale bar = 500µM), 200x magnification (ii; Scale bar = 100µM) and staining (iii, Scale bar as shown) for PanCK (Green), CD163 (Yellow), CD8 (Red) and DNA (Blue).

**Table 1.** Clinicopathological parameters of the study cohort. ER+: Estrogen receptor positive, HR: Hormone receptor, HER: human epidermal growth factor receptor 2, LVI: Lymphovascular invasion. BMI: Body Mass Index

|  | Level | Overall |
|---|---|---|
| n |  | 43 |
| Age at Diagnosis [Mean(SD)] |  | 54.02 (13.59) |
| BMI [Mean(SD)] |  | 33.18 (7.04) |
| Race (%) | White | 24 (55.8) |
|  | Black or African American | 18 (41.9) |
|  | Unknown | 1 (2.3) |
| Ethnicity (%) | Not Hispanic or Latino | 24 (55.8) |
|  | Hispanic or Latino | 1 (2.3) |
|  | Unknown | 18 (41.9) |
| Menopausal Status at Diagnosis (%) | Post-menopause | 27 (62.8) |
|  | Pre-menopause | 14 (32.6) |
|  | Pregnant/Breast Feeding | 2 (4.7) |
| ER (%) | Positive (≥ 10) | 25 (58.1) |
|  | Low positive (1- < 10) | 3 (7.0) |
|  | Negative (0- < 1) | 15 (34.9) |
| PR (%) | Positive (≥ 10) | 15 (34.9) |
|  | Low positive (1- < 10) | 8 (18.6) |
|  | Negative (0- < 1) | 20 (46.5) |
| HER2 (%) | Positive | 6 (14.0) |
|  | Equivocal | 1 (2.3) |
|  | Negative | 36 (83.7) |
| Histology (%) | Invasive Ductal | 35 (81.4) |
|  | Invasive Lobular | 2 (4.7) |
|  | Invasive Micropapillary | 2 (4.7) |
|  | Invasive Mixed Ductal and Lobular | 3 (7.0) |
|  | Invasive Other | 1 (2.3) |
| Clinical Stage (%) | IIIB | 12 (27.9) |
|  | IIIC | 15 (34.9) |
|  | IV | 16 (37.2) |
| Primary Tumor T (%) | T4d | 43 (100.0) |
| Regional Lymph Nodes N (%) | N1 | 14 (32.6) |
|  | N2 | 1 (2.3) |
|  | N2a | 1 (2.3) |
|  | N3 | 3 (7.0) |
|  | N3a | 6 (14.0) |
|  | N3b | 3 (7.0) |
|  | N3c | 15 (34.9) |
|  | M0 | 27 (62.8) |
| Distant Metastases M at initial diagnosis (%) | M1 | 16 (37.2) |
| Pathological Staging Tumor ypT (%) | T0 | 1 (2.3) |
|  | T1b | 1 (2.3) |
|  | T1c | 1 (2.3) |
|  | T1mic | 1 (2.3) |
|  | T2 | 7 (16.3) |
|  | T3 | 13 (30.2) |
|  | T4b | 2 (4.7) |
|  | T4d | 17 (39.5) |
| Pathological Staging Node ypN | N0 | 2 (4.7) |
|  | N0(i-) | 1 (2.3) |
|  | N1a | 1 (2.3) |
|  | N2 | 6 (14.0) |
|  | N2a | 11 (25.6) |
|  | N3 | 3 (7.0) |
|  | N3a | 19 (44.2) |
| Number of Nodes Removed [Mean(SD)] |  | 21.05 (9.73) |

Unsupervised hierarchical clustering across data from all regions and cases demonstrated segregation of adjacent normal epithelial regions from malignant compartments, whereas tumor stroma, leading edge, and LVI did not segregate into obvious clusters (**Figure 2A**). The first two key components in a principal component (PC) analysis captured a relatively small proportion of cumulative variance: 16.2% for PC1 and 6.3% for PC2, with the normal-malignant status separating along the PC1 axis (**Figure 2B**). When examining principal component loadings, the *keratins 5, 14*, *15,* and *17* correlated along PC1 in the positive direction. *COL3A1* correlated in the negative direction (**Figure 2C**). These loadings were consistent with differential gene expression analysis.

**Figure 2.**
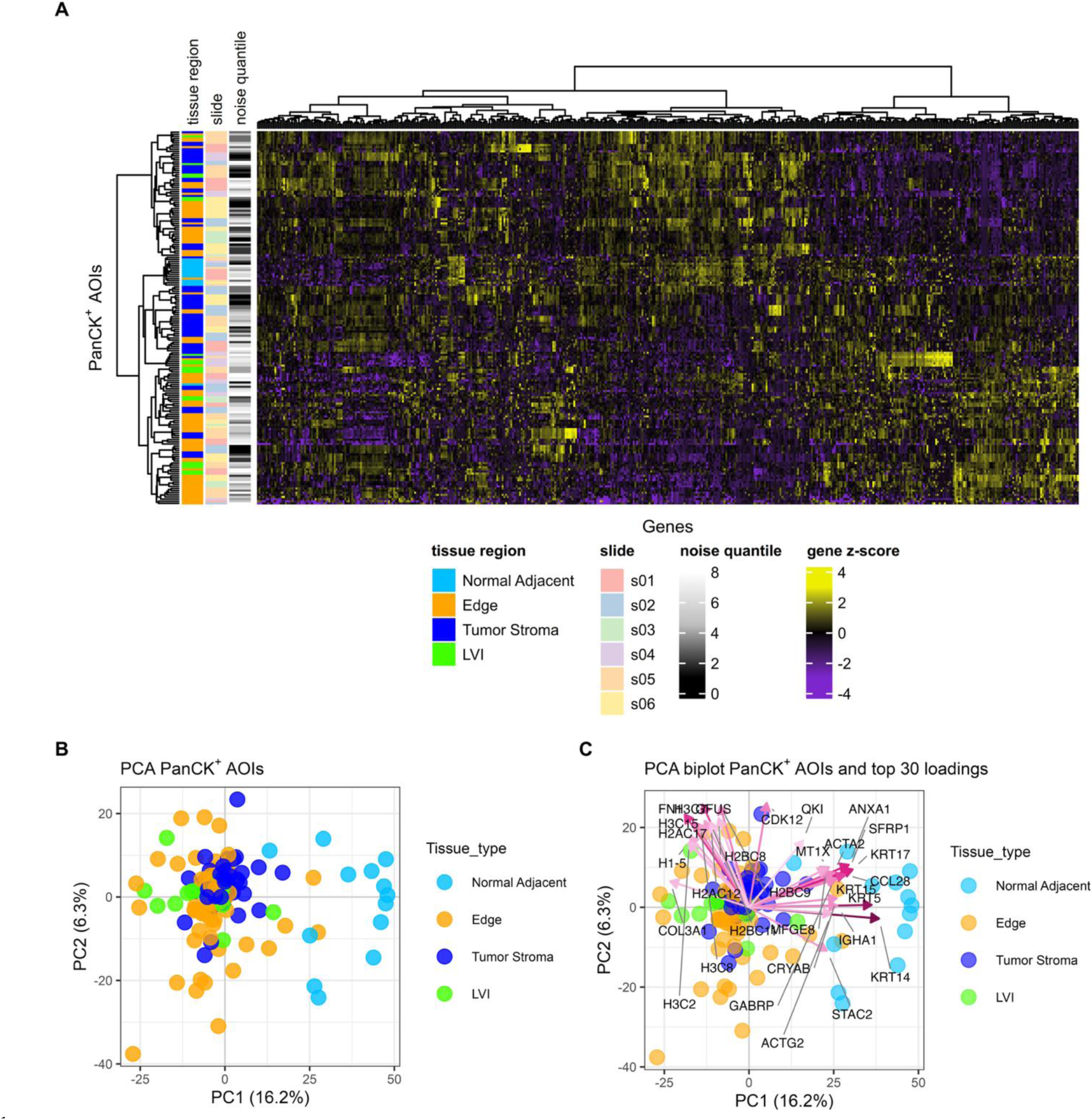
Spatial transcriptomic profiling identifies distinct epithelial states in malignant and normal adjacent tissue compartments. **(A)** Heat map of Highly Variable Genes (HVG) with unsupervised clustering showing tissue segment, slide, and noise quantiles. Each row represents a PanCK+ area of illumination (AOI), which after quality control yielded 84 tumor edge, 65 stromal tumor, 19 LVI, and 15 normal adjacent tissue segments. **(B)** Principal component (PC) plots of spatial transcriptomic data batch corrected for slides and patients colored by annotated tissue region. (C) Principal component loadings (top 30 shown) of the first two PCs shown in A.

When comparing tumor stroma, leading edge, and LVI compartments, respectively, to adjacent normal tissue, all three compartments showed 5-fold or greater downregulation of *keratins 5* and *14* and greater than 5-fold upregulation of *COL3A1* (p_all_ < 0.05). A total of 182 genes showed 5-fold or greater downregulation across all three tumor compartments compared to adjacent normal tissue, and 90 genes showed 5-fold or greater upregulation across all tumor compartments (**Figure 3A, 3B, 3C** and **Supplementary Table 1**). We then examined differential gene expression between the respective tumor compartments. Consistent with the PCA plot showing similar clustering of tumor regions, between tumor stroma and tumor leading edge *CD24* was the only significantly upregulated gene in tumor middle compared to leading edge (**Figure 3D**), *IFITM1*, *PARP12*, *HLA-F*, and *RAD21* were the only significantly upregulated genes in LVI regions when comparing between tumor stroma and LVI regions (**Figure 3E**), and there were no differentially expressed genes when comparing between tumor leading edge and LVI regions (**Figure 3F**). Overall, these data support known conserved differences between cancerous and normal states. However, because gene pathway analyses are sensitive to modest, correlated gene expression changes^33^, we hypothesized that individual tumor compartments would exhibit conserved malignant programs compared to normal tissues, and unique transcriptional programs when compared to each other.

**Figure 3.**
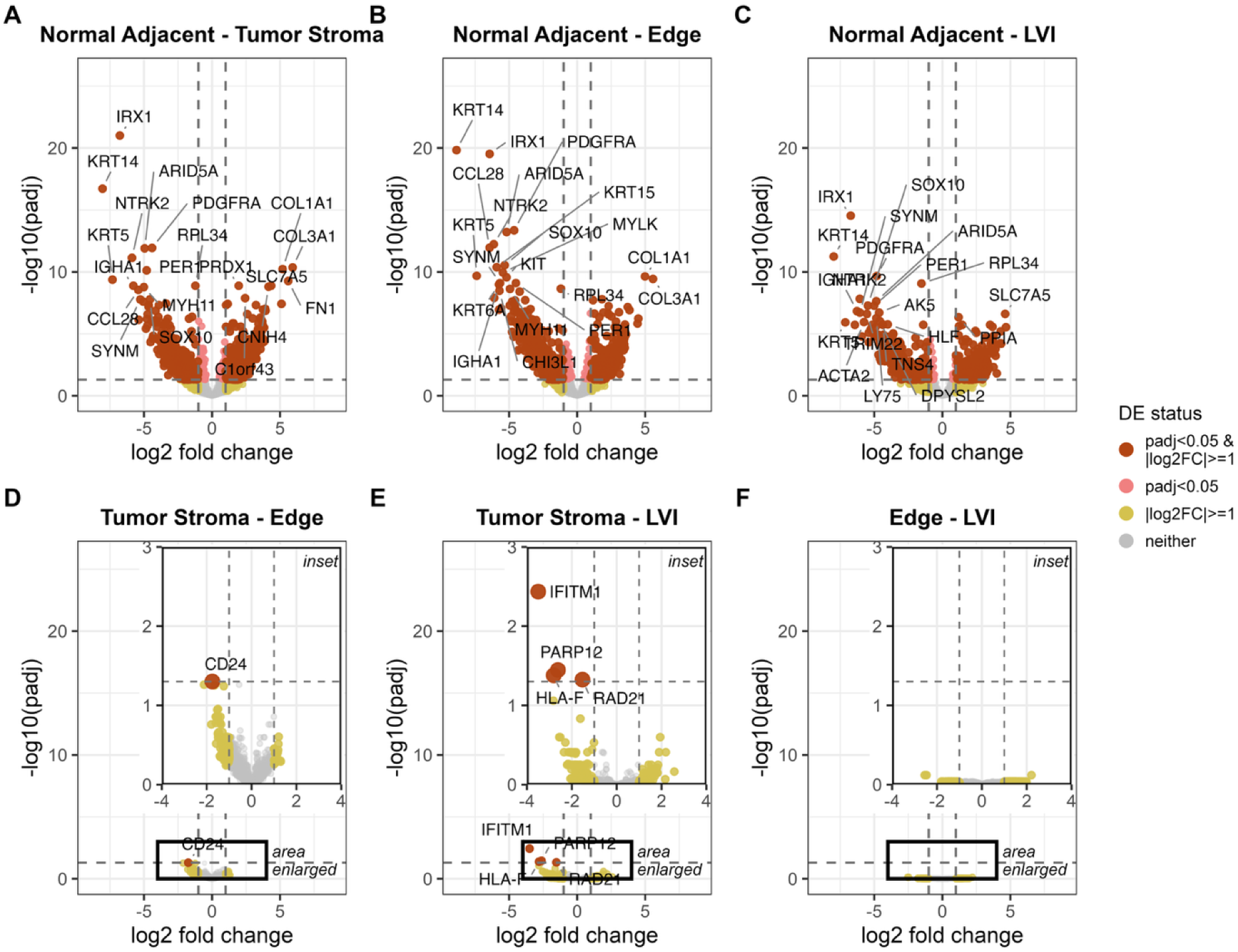
Differential gene expression between adjacent normal and tumor compartments. (**A-F**) Volcano plots of differentially expressed (DE) genes (red points, padj < 0.05 and |log2fc >=1|) for PanCK+ segment AOIs. Positive log2 fold change indicates higher expression in the right-listed segments of each contrast. Differential expression was fit in limma with fixed effects for tissue and slide. Within-patient correlation was estimated with duplicateCorrelation (block = Patient; consensus correlation = 0.2414238) and supplied to lmFit. Pairwise tissue contrasts were moderated with empirical Bayes (eBayes, trend = TRUE) and Benjamini–Hochberg-adjusted.

### Spatial Context Reveals Differences in Adjacent Normal and Tumor Compartment Hallmarks of Cancer

To define conserved and compartment-specific malignant transcriptional programs, we performed gene set enrichment analysis (GSEA) using the Hallmarks of Cancer from the Molecular Signatures Database (MSigDB), comparing each tumor compartment with adjacent normal tissue and with the other tumor compartments^34^.

Relative to adjacent normal tissue, tumor stroma, leading edge, and lymphovascular invasion (LVI) compartments shared a malignant transcriptional program, with 14 Hallmark pathways significantly enriched across tumor compartments, including E2F Targets, MYC Targets, mTORC1 Signaling, and Reactive Oxygen Species (**Figure 4A**). Conversely, 12 pathways were commonly suppressed relative to adjacent normal tissue, including TNFα Signaling via NF-κB, Apical Junction, and p53 Pathway. Thus, despite their distinct spatial locations, malignant compartments exhibited a conserved transcriptional state relative to normal regions.

**Figure 4.**
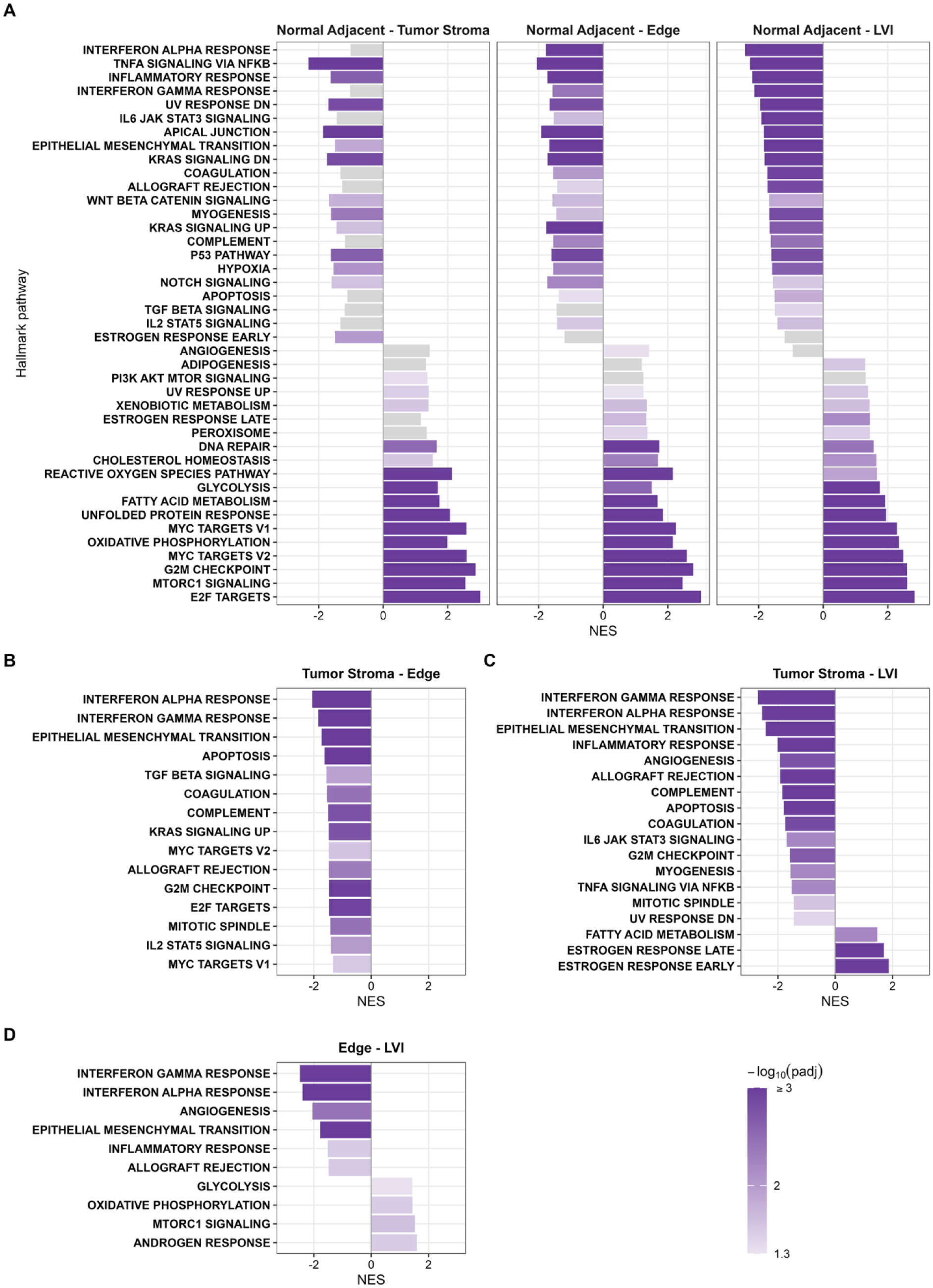
Spatial context reveals differences in adjacent normal and tumor compartment Hallmarks of Cancer. Gene Set Enrichment Analysis (GSEA) of the Molecular Signatures Database Hallmark of Cancer Pathways shows gene set enrichment sorted by the enrichment scores of tumor regions compared to normal (**A**). This analysis was repeated for tumor stroma and leading edge (**B**), tumor stroma and LVI (**C**), and then tumor leading edge and LVI (**D**). Bars are colored by adjusted *P* value. Gray bars are not significant.

In contrast, direct comparison of tumor compartments revealed pronounced spatial divergence. Relative to tumor stroma, leading edge regions showed broad suppression of Hallmark pathways, most prominently Interferon Alpha Response, Interferon Gamma Response, and Epithelial-Mesenchymal Transition (EMT) (**Figure 4B**). LVI exhibited a similar suppression of immune-associated programs relative to tumor stroma, with 15 pathways downregulated, including Interferon Alpha and Gamma Responses and EMT. Notably, LVI uniquely exhibited enrichment of Early and Late Estrogen Response and Fatty Acid Metabolism (**Figure 4C**). Compared with the leading edge, LVI further showed reduced Interferon Alpha and Gamma Responses, Inflammatory Response, Angiogenesis, and EMT, together with increased Androgen Response, mTORC1 Signaling, and Oxidative Phosphorylation (**Figure 4D**).Together, these data reveal a spatially organized transcriptional adaptation across tumor compartments wherein the IBC tumor cells retain core malignant programs across compartments but acquire distinct states within the tumor microenvironment. In particular, the LVI compartment is characterized by attenuation of immune and inflammatory programs and increased hormone-responsive and metabolic signaling, consistent with adaptation to the lymphovascular niche. We next used Gene Ontology analysis to define the biological processes, molecular functions, and cellular components underlying these compartment-specific states. Complete Hallmark pathway results are provided in **Supplementary Table 2**.

### IBC Tumor Compartments Exhibit Distinct Immune, Structural, and Metabolic States

Gene ontology analysis comparing the leading edge of the tumor compared to tumor middle showed broad suppression of processes associated with cellular organization and immune response, including cell adhesion, actin cytoskeleton, keratinization, intermediate filament organization, cellular matrix adhesion and structural components, collagen binding, T cell activation, and humoral immune response compared to tumor stroma (**Figure 5A, B, C Supplementary Table 3, 4, 5**). No biological process, molecular function, or cellular component was significantly enriched in the leading edge relative to tumor stroma, suggesting that this compartment was characterized predominantly by loss of structural and immune programs rather than acquisition of a distinct transcriptional program.

**Figure 5.**
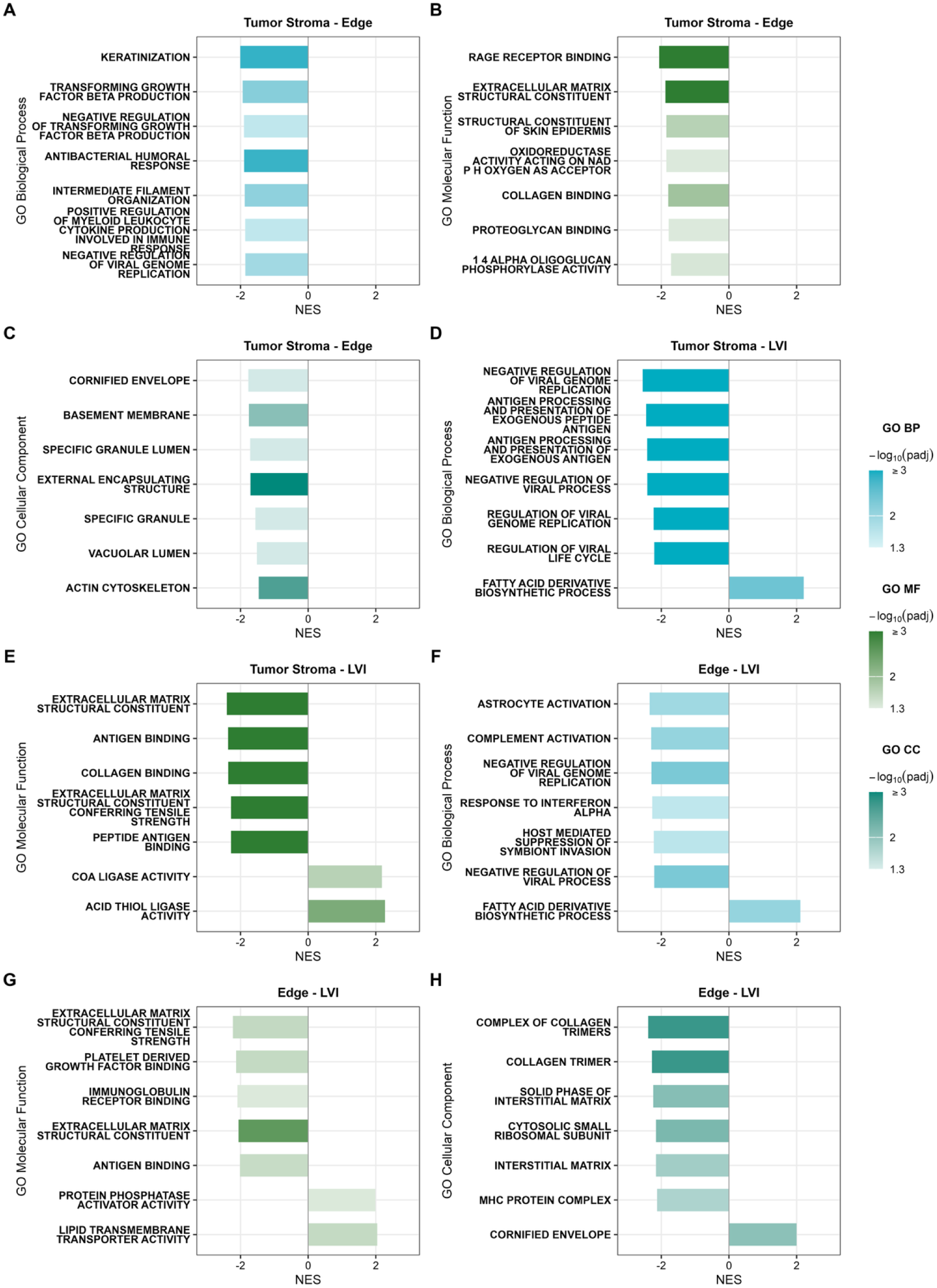
IBC Tumor Compartments Exhibit Distinct Immune, Structural, and Metabolic States. Gene Ontology analysis when comparing tumor stroma and leading-edge biologic processes (**A**), molecular functions (**B**), and cellular components (**C**). This was repeated for tumor stroma and LVI biologic processes (**D**), molecular functions (**E**), and then tumor leading edge and LVI biologic processes (**F**), molecular functions (**G**), and cellular components (**H**).

In contrast, LVI when compared to tumor middle was characterized by reduced programs involved in antiviral responses, antigen processing and presentation, and regulation of the immune response, including decreased antigen and peptide binding and reduced MHC protein complex representation (**Figures 5D, E; Supplementary Tables 3,5**). Concurrently, LVI showed enrichment of lipid metabolic programs, including fatty acid derivative biosynthesis, cholesterol biosynthesis through desmosterol, and acyl-CoA metabolism. These findings identify the LVI compartment as a metabolically distinct niche characterized by attenuation of immune surveillance and increased lipid metabolic activity.

Comparison of LVI with the tumor leading edge further supported this compartmental distinction. LVI exhibited reduced antiviral and interferon-α response programs, together with increased fatty acid derivative biosynthesis and lipid transmembrane transporter activity (**Figures 5F, G, H; Supplementary Tables 3–5**). LVI also showed reduced extracellular matrix and platelet-derived growth factor binding programs and increased representation of mitochondrial matrix and cornified envelope components. Thus, although both LVI and the tumor leading edge displayed reduced immune-associated programs, LVI was distinguished by a stronger metabolic and lipid-associated transcriptional state.

Most importantly, the tumor stroma showed notable upregulation in biological processes, molecular functions, and cellular components related to cellular division, dedifferentiation, and hypoxia compared to both LVI regions and tumor leading edge regions (**Supplementary Tables 3, 4,** and **5**).

Together, these findings reveal that IBC malignant regions are not transcriptionally uniform but instead occupy distinct spatial states wherein the tumor stroma is enriched for proliferative, hypoxic, and dedifferentiated programs, the tumor leading edge shows broad loss of structural and immune programs, and LVI is characterized by immune attenuation coupled to lipid metabolic activation.

### Tumor-Lymphatic Biomimetic Model Recapitulates Transcriptomic State of Human LVI

We recently developed a bioengineered platform (T-LAB) that recapitulates key fluidic topographical features of the tumor lymphatic interface and enables modeling of collective invasion and lymphatic dissemination^32^. To determine whether T-LAB recapitulates features of the human IBC lymphovascular invasion (LVI) niche, the well-characterized, patient-derived, treatment-naïve IBC cell line SUM149 was introduced into the central chamber and allowed to invade through the extracellular matrix into a lymphatic vessel-like compartment. Tumor cells recovered from the lymphatic compartment were designated T-LAB circulating cells (TLAB-CC) (**Figure 6A**).

**Figure 6.**
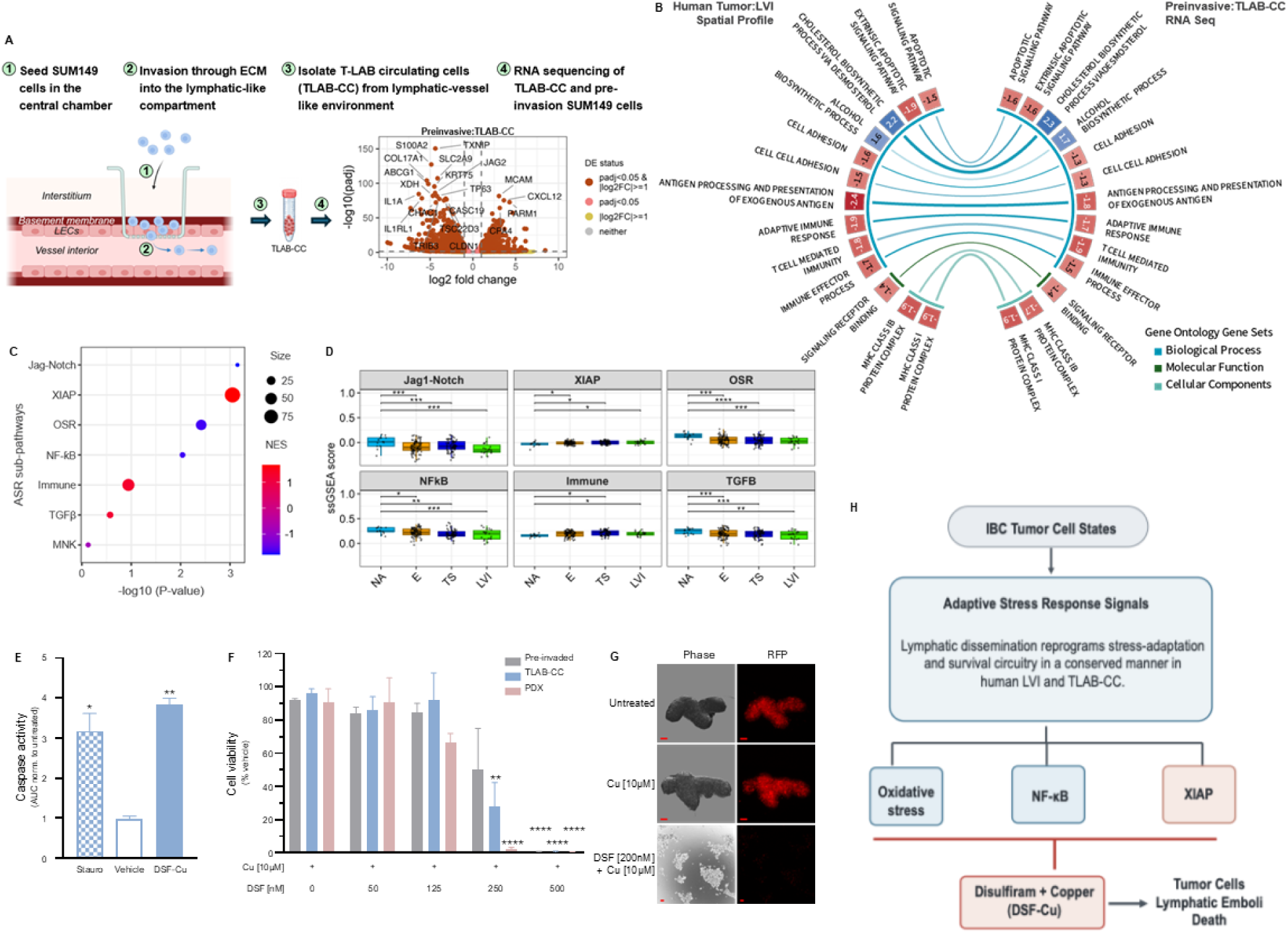
A Biomimetic T-LAB Model Recapitulates Human IBC LVI and Reveals a Pharmacologically Actionable Vulnerability. **(A)** Steps in isolation of T-LAB circulating cells and RNA sequencing resulting in volcano plots of differentially expressed (DE) genes (red points, padj < 0.05 and |log2fc >=1|) for TLAB-CC vs Preinvasive SUM149 cells. Positive log2 fold change indicates higher expression in the TLAB-CC. **(B)** CIRCOS plot showing shared significant pathway enrichment in IBC human specimen and the preclinical model. **(C)** Bubble plot showing significant gene enrichment for gene sets related to the ASR sub-pathways in TLAB-CC vs preinvasive SUM149. Bubble size shows the number of genes present for each pathway. (**D)** Single-sample Gene Set Enrichment Analysis (ssGSEA) analysis for the ASR sub-pathways in Normal Adjacent (NA), Edge (E), Tumor Stroma (TS) and LVI regions where each ROI is independently scored based on the expression of genes in each gene set identifies up-regulated ASR sub-pathway (XIAP and Immune) and down-regulated ASR sub-pathway(OSR, Jag1-Notch, NFƙB and TGFβ) with red bullets showing enrichment in TLAB-CC and blue bullets showing downregulation in TLAB-CC. (**E)** Caspase activity measured in cells following treatment with positive control (Staurosporine/Stauro;10µM), vehicle, or DSF [500nM] in presence of Cu [10µM], as area under the curve (AUC) for 22h normalized to untreated and compared with the control. Bars represent mean ± SEM [n=2 (6)]. Statistical significance was determined using Tukey test. *P*-value is represented as *< 0.05, \**\** < 0.01, \*\**\** < 0.001, \**\*\*\** < 0.0001 when compared with vehicle. (**F)** Dose-dependent measurement of viability in preinvasive parental (SUM149; grey bar), TLAB-CC (Blue bar) and MDA-IBC3 (Pink bar) treated with DSF+Cu (as shown) at 24H. Cell viability was determined by cell titer glow assay (Promega) where cell viability was calculated as luminescence with treatment/ luminescence with vehicle control (highest concentration of DMSO) ×100. n = 3(3). Bars represent mean ± SEM. Statistical significance was determined using Tukey test. *P*-value is represented as *< 0.05, \**\** < 0.01, \*\**\** < 0.001, \**\*\*\** < 0.0001 when compared with Cu condition for the specific cell line. (**G)** Effect of pharmacological inhibitor, DSF-Cu on formation of tumor emboli derived from TLAB-CC; Representative images at clinically relevant doses of the drug at 24h post treatments are shown at the top of each column. Bar represents 100µm. (**H)** Integrated model identifies a conserved stress response program across human tumor niches for mechanistic testing and drug discovery.

We performed whole-transcriptome RNA sequencing of TLAB-CC and matched pre-invasion SUM149 cells to define transcriptional changes associated with lymphatic dissemination (**Figure 6A** volcano plot**; Supplementary Table 6**). TLAB-CC exhibited extensive transcriptional remodeling, with 2,560 upregulated and 2,543 downregulated relative to pre-invasion cells.

Gene ontology analysis comparing TLAB-CC relative to pre-invasion cells revealed selective conservation of biological programs identified in human IBC LVI by spatial transcriptomic profiling. Both TLAB-CC and human IBC LVI showed reduced programs related to apoptotic signaling, cholesterol biosynthesis, and cell adhesion, while immune-associated programs involving antigen processing and presentation, exogenous antigen presentation, adaptive immune responses, and T cell–mediated immunity were also suppressed in both compartments (**Figure 6B**). Of the 207 biological processes dysregulated in human LVI, 74 (37%) were concordantly dysregulated in TLAB-CC (**Supplementary Table 6**). In contrast, concordance was limited at the molecular-function and cellular-component levels, with 1 of 28 and 2 of 23 pathways, respectively, showing concordant dysregulation (**Supplementary Table 6**). Notably, both concordant cellular-component pathways involved MHC class I protein complexes. Although T-LAB did not globally reproduce the human LVI transcriptome, it notably recapitulated biological programs associated with immune regulation, cell adhesion, survival, and metabolism. Together, these findings establish TLAB-CC as a biomimetic model that captures key biological features of the human LVI transcriptional state providing a system for mechanistic interrogation of tumor-cell adaptation within the lymphatic niche.

### Adaptive Stress Response Is Conserved Across Human IBC LVI and T-LAB-Derived Tumor Cells

Lymphovascular dissemination exposes tumor cells to diverse microenvironmental stresses, including oxidative, inflammatory, and immune stress. We therefore asked whether the transcriptional state acquired during lymphatic dissemination reflects a conserved adaptive stress response (ASR) program. We previously defined an ASR-226 metagene and associated signaling pathways implicated in tumor adaptation to microenvironmental stress^16,29,31^. Given the shared survival and immune-associated programs identified in human LVI and TLAB-CC (**Figure 6B**), we examined the ASR framework across both systems.

TLAB-CC exhibited coordinated remodeling of multiple ASR-associated pathways, including significant suppression of JAG1–Notch, oxidative stress response, and NF-κB programs. XIAP signaling was significantly increased. Immune and TGF-β signaling showed trends toward increased activity, whereas MNK signaling showed a trend toward suppression (**Figure 6C**). Importantly, human LVI exhibited a similar pattern, with suppression of JAG1–Notch, oxidative stress response, TGF-β, and NF-κB programs and increased XIAP and immune-associated signaling (**Figure 6D**).

### T-LAB Reveals a Pharmacologically Actionable Vulnerability Associated with Lymphatic Dissemination

We next asked whether tumor cells acquiring the LVI-associated state retained therapeutically actionable vulnerabilities. Because prior studies have demonstrated disulfiram/copper (DSF-Cu) negatively impacts lipid metabolism, a vulnerability we identified in LVI regions, while also inducing oxidative stress, a vulnerability we identified in stromal regions, we used disulfiram/copper (DSF-Cu) as a proof-of-principle pharmacologic perturbation to examine if it would successfully result in apoptotic cell death of TLAB-CC (**Figure 6E**) cells representing stromal and LVI tumor regions^28,35,36^; DSF-Cu reduced the viability of both parental SUM149 cells and TLAB-CC in a dose-dependent manner, with TLAB-CC showing greater loss of viability at the intermediate 250-nM disulfiram dose (**Figure 6F**). At 500 nM, DSF-Cu reduced viability to near-background levels in both populations. Interestingly, in a separate IBC PDX (MDA-IBC3) DSF-Cu induced cell death and apoptosis at near similar levels to SUM149 and TLAB-CC indicating our human based data is recapitulated in multiple IBC models (**Figure 6F**). Moreover, targeting OSR pathway with DSF+Cu also inhibited TLAB-CC tumor emboli formation in a dose dependent manner and 200nM DSF in presence of 10µM Cu, induced cell death in the *in vitro* tumor emboli model of TLAB-CC (**Figure 6G**). These findings indicate that acquisition of the LVI-associated transcriptional state does not eliminate oxidative-stress vulnerability and may increase sensitivity to copper-dependent pharmacologic stress. Furthermore, the T-LAB platform provides a functional model for interrogating therapeutic vulnerabilities associated with lymphatic dissemination. To demonstrate this utility, SUM149 cells cultured on the T-LAB platform³² were treated with 250 nM DSF and 10 µM Cu resulted in a marked reduction in cell viability by Day 3, with only ∼1.25% viable cells remaining in the trans-well and no detectable cells in the lymph-like fluidic compartment. These findings suggest that DSF-Cu treatment effectively inhibits tumor cell invasion through the T-LAB platform (Data not shown), compared with the invasion observed in our previous study³².

The concordance of ASR-associated pathway remodeling between human LVI and TLAB-CC indicates that lymphatic dissemination is accompanied not by uniform activation of stress pathways, but by selective reprogramming of stress-adaptation and survival circuitry. Notably, the sensitivity of TLAB-CC to copper-dependent oxidative stress occurs in the context of this coordinated remodeling of oxidative stress, NF-κB, and XIAP signaling, linking the transcriptional state of LVI-like tumor cells to a pharmacologically actionable stress vulnerability. Together, these findings identify ASR as a conserved transcriptional framework (**Figure 6F**) linking the human LVI niche to a biomimetic lymphatic dissemination model and establish T-LAB as a tractable system for functional interrogation of stress-adaptation mechanisms that may support tumor-cell survival during lymphatic dissemination.

## Discussion

Inflammatory breast cancer is an aggressive disease characterized by rapid progression, extensive dermal lymphatic invasion, and early metastatic spread. While these features are well recognized, the tumor cell adaptations that enable this process, especially within the lymphovascular compartment, have remained poorly understood. In this study, using spatially resolved whole-transcriptome profiling of human IBC specimens, we show that tumor cells share core transcriptional programs but also adopt distinct, context-dependent states across tumor stroma, leading edge, and lymphovascular regions. These findings suggest that tumor cells are not transcriptionally distinct clones but rather dynamically adapt across niches in their local microenvironment that facilitate their survival, immune evasion, and dissemination. We further demonstrated in a translational model that copper disulfiram can kill tumor cells associated with metastasis as well as those confined to the stroma suggesting that therapeutics targeting pathways across tumor niches may offer improved disease control.

A central insight from this work is the emergence of a spatially organized continuum of tumor cell states that appears to track with the process of dissemination. Tumor stroma regions are enriched for proliferative and survival programs, the leading edge adopts features consistent with invasion including reduced cell-cell adhesion and cytoskeletal remodeling, and LVI regions display the strongest signatures of immune evasion alongside metabolic adaptation to fatty acid utilization. Together, these observations support a directional model in which tumor cells transition from a proliferative stroma state to an invasive leading-edge phenotype and ultimately to a lymphovascular-adapted state optimized for survival within the lymphatic niche (stroma to leading edge to LVI).

Although LVI is a well-established predictor of poor outcomes across cancers^10,12,37–40^, its biology has been difficult to study due to the small size and limited cellularity of these lesions. Here, we provide what is, to our knowledge, the first spatial transcriptomic analysis focused specifically on tumor cells within IBC LVI regions. Remarkably, we observed relatively few differences at the level of individual genes when comparing LVI to other tumor compartments. However, pathway-level analyses revealed a clearer signal wherein LVI tumor cells showed strong enrichment of immune evasion programs along with a shift toward fatty acid metabolism compared to the tumor stroma or leading edge. This is consistent with the lipid-rich composition of lymphatic fluid and aligns with prior work linking impaired interferon signaling^41–43^ and fatty acid oxidation^44^ to metastatic progression. These findings suggest that tumor cells entering lymphovascular spaces adopt a state that is both immunologically evasive and metabolically suited to this environment. These features highlight potential therapeutic opportunities, including restoring immune recognition within LVI niches or targeting lipid metabolic dependencies.

Another core finding of our study was that tumor cells within the leading edge had a distinct set of adaptations consistent with local invasion. These regions were characterized by reduced cell–cell adhesion, cytoskeletal reorganization, and enrichment of epithelial–mesenchymal transition (EMT)-related pathways. These findings are consistent with a role for EMT in facilitating local invasion and are corroborated by a spatial study in squamous carcinomas, which found leading edge regions are characterized by alterations in adhesive signaling, as well as others who have shown cytoskeletal rearrangement is core to tumor cell metastasis^45^. Interestingly, the leading edge of the tumor demonstrated increased pathways related to EMT compared to LVI regions which is consistent with models demonstrating successful metastatic colonization requires reversal of EMT through mesenchymal-epithelial transition (MET).^46,47^ Our results support this dynamic model in which tumor cells transition between phenotypic states as they move from invasion into intravascular survival.

Tumor stroma regions, in contrast, appeared to represent a more proliferative and canonical tumor state. These regions showed strong enrichment of pathways associated with cell division and survival, including KRAS signaling, consistent with prior bulk sequencing studies in breast cancer and IBC^48–50^. Notably, many hallmark cancer pathways were most prominent in the tumor stroma compared to the tumor leading edge and tumor LVI regions, highlighting how bulk transcriptomic approaches may be dominated by this majority compartment and may overlook critical adaptations occurring in less abundant but functionally important regions such as the leading edge and LVI that are revealed by digital spatial profiling.

Finally, our findings demonstrate that the T-LAB bioengineered platform^32^ recapitulates key transcriptional features of human IBC LVI, providing a tractable model for investigating tumor-cell adaptation during dissemination. TLAB-derived circulating cells exhibited alterations in cholesterol and alcohol metabolism, as well as reduced antigen processing and presentation, including downregulation of genes associated with MHC-class 1 complexes. These changes were also observed in human IBC LVI regions, supporting the biological relevance of the model while highlighting metabolic remodeling and altered antigen presentation as features of the dissemination-associated state. Importantly, T-LAB biomimetic enables functional interrogation of vulnerabilities shared across distinct tumor-cell states. As proof of principle, disulfiram-copper treatment reduced the viability of both preinvasive, stromal-like tumor cells and TLAB-CC, suggesting that pharmacologic targeting may affect tumor cells across spatially distinct states. Together, these findings establish T-LAB as a reproducible in vitro platform for investigating tumor-intrinsic mechanisms associated with invasion, lymphatic dissemination, and potential immune evasion, and for evaluating therapeutic strategies against vulnerabilities that persist across tumor-cell states.

While the distinct transcriptional programs identified in this study represent a major advance in understanding IBC biology, there are several limitations to the current study. The rarity of IBC and the small size of LVI lesions limited the number of regions available for analysis, preventing robust correlations with clinical outcomes. In addition, because analyses were aggregated across patients, some findings may reflect interpatient variability. Finally, while our spatial approach enabled detailed transcriptional characterization, functional validation of these pathways remains an important next step. Despite these limitations, this study provides a framework for understanding how tumor cells adapt across spatial niches within human IBC. We identify a distinct lymphovascular state characterized by reduced antigen presentation, dampened inflammatory signaling, and increased reliance on lipid metabolism. More broadly, our findings support a model in which metastatic competence is not a fixed property of tumor cells but instead emerges through dynamic, microenvironment-driven adaptation along a spatial continuum. By defining these compartment-specific states directly in human tumors, this work highlights potential vulnerabilities including fatty acid metabolism in LVI regions, cytoskeletal remodeling at the leading edge, and proliferative signaling in tumor stroma, which could be targeted to interfere with metastatic progression in inflammatory breast cancer.

## Declaration of Generative AI and AI-assisted technologies

During the preparation of this work, the authors used copilot and ChatGPT to assist with grammatical edits of the manuscript. After using this tool/service, the author(s) reviewed and edited the content as needed and take full responsibility for the content of the published article. Figures 2 through 6 are created directly from the underlying data in Rstudio 4.4.1 with code writing and formatting assisted by Claude Opus 4.8.

## Data and Code availability

The code and datasets related to gene expression used in this manuscript are available at https://github.com/The-Devi-Lab without restrictions. Any other datasets used in this manuscript are available from the corresponding author on reasonable request.

## Author contributions

G.R.D. D.P.M. W.A.W. - conceptualization, methodology, analysis, resources, supervision. D.P.M., G.R.D., P.P. - original draft preparation, writing, reviewing and editing, supervision. J.D. - molecular pathology, conceptualization, histology. I.F. - pathology. S.K, J.N. - tissue microarray design, pathologic assessment. D.P.M. S.G., A.F. - bioinformatic analysis. D.P.M. P.P., S.G. - methodology, analysis, validation, writing, editing, reviewing. All authors-reviewing, editing. All authors have read and agreed to the published version of the manuscript.

## Acknowledgements

The authors thank Duke Biorepository and Precision Pathology Center team: Xufeng Cheng, Parker Shoaf; Katherine Frankey; Dr. Kouros Owzar and Dr. Yanwan Dai at the Duke Cancer Institute Biostatistics and Bioinformatics for technical support. The Hollings Cancer Center CURE HUB: Cancer Research and Education Hub for Unified Biomedical, Clinical, and Population Science team: Zara Meaby, Hesham El-Shewey, Dr. William G. Hawkins for technical support and/or helpful discussions.

## Methods

### Specimen procurement

Tumor Microarray (TMA) in triplicate of 43 IBC surgical samples obtained after standard of care neoadjuvant systemic therapy from confirmed cases enrolled on the prospective IBC registry from MD Anderson Cancer Center Morgan Welch Inflammatory Breast Cancer Clinic and Research Program (ClinicalTrials.gov Identifier: NCT02876107, registered on August 22, 2016)^51^. Informed consent was obtained from all subjects and/or their legal guardian(s). IBC specialists reviewed all cases in a prospective manner. Overall, patient inclusion criteria include histological confirmation of breast carcinoma, IBC confirmed according to international consensus criteria^3^. All experimental protocols were approved by the institutional committees, and all methods were carried out in accordance with relevant guidelines and regulations.

### Digital Spatial Profiling (DSP) of RNA

RNA profiling using Digital Spatial Profiling (DSP) was conducted at Duke University with the Whole Transcriptome Atlas kit on the NanoString GeoMx Digital Spatial Profiler. Tissue sections were treated with oligonucleotide probes along with fluorescently labeled antibodies targeting epithelial cells (PanCK, Catalogue #121300310, Nanostring), Macrophage (CD163, Catalogue #ab272103, Abcam clone EPR14643-36), and cytotoxic T cells (CD8, Catalogue #AB196193, Abcam clone EP115Y). Regions of interest (ROIs) were carefully delineated to include clusters of cells with similar histological characteristics, ensuring that no ROI exceeded the 700 μm² area limit. Within each ROI, segments stained with different antibodies were classified as areas of illumination (AOIs). Library preparation was done following the protocols outlined in the NanoString GeoMx Library Preparation Manual. Libraries were then normalized to equimolar concentrations and sequenced using the Illumina NovaSeq 6000 platform, targeting a depth of 30 read pairs per μm². Data processing, including barcode trimming, alignment, and deduplication based on unique molecular identifiers (UMIs), was carried out on the GeoMx DSP Analysis Server. The resulting UMI count matrix for each AOI was subsequently analyzed using R/Bioconductor^52^.

### Quality control and normalization

Quality control (QC) assessments were carried out for each area of illumination (AOI), evaluating metrics such as signal-to-noise area under the curve (AUC) and the limit of detection (LOD), following established methods^53–55^. AOIs were excluded from downstream analysis if they failed to meet the following thresholds: (i) a minimum of 15,000 unique molecular identifier (UMI) counts, (ii) a signal-to-noise AUC greater than 0.55, and (iii) more than 20% of genes detected above the LOD. Additionally, genes that fell below the LOD in over 75% of AOIs were filtered out. To address technical variability and background noise, background subtraction and quantile normalization were applied as described previously ^55,56^.

### Dimension reduction and clustering

Highly variable genes (HVGs) were identified from the slide batch-corrected log₂-normalized CPM matrix. Within each slide, a local polynomial regression (LOESS, span = 0.25) was fit to the gene-level mean–variance relationship with a variance floor of 1%. Genes were then ranked within each slide by the ratio of observed to LOESS-predicted variance. A gene was retained as an HVG if it ranked in the top 150 of at least one slide and in the top 5,000 of the dataset-wide LOESS-rank. HVG values were z-scored by gene and rendered as a heatmap with ComplexHeatmap^57^. Clustering was done independently on rows and columns with Ward.D2 hierarchical clustering of Euclidean distance. Principal component analysis was computed on the AOI × HVG matrix after residualizing slide and patient effects with limma::removeBatchEffect and preserving tissue identity in the design^58^.

### Differential expression analysis

Differential expression (DE) analysis was performed using the limma package^58^ (RRID: SCR_010943). For each gene, a linear model was fit with fixed effects for tissue zone and slide. Multiple AOIs per patient were accommodated by estimating the within-patient correlation with duplicateCorrelation() (block = Patient; consensus correlation = 0.2414238) and supplying that value to lmFit(). Six pairwise tissue contrasts (middle vs normal, edge vs normal, LVI vs normal, edge vs middle, LVI vs middle, and LVI vs edge) were extracted with contrasts.fit() and moderated by empirical Bayes (eBayes(), trend = TRUE). Genes were considered differentially expressed if they showed an absolute log2 fold change greater than 1.0 and Benjamini-Hochberg adjusted q-value < 0.05. Heatmaps were created using the pheatmap package in R^57^, with hierarchical clustering of both rows and columns carried out using the Ward.D2 linkage on Euclidean distance.

### Gene set enrichment analysis

Gene set enrichment analysis (GSEA) was carried out using the fgsea package (RRID: SCR_020938) for six pairwise tissue contrasts. Genes were ranked within each contrast based on their signed log₂ fold change from the limma fit and tested against the MSigDB Hallmark collection and the three Gene Ontology categories (Biological Process, Cellular Component, and Molecular Function). Pathways with Benjamini–Hochberg-adjusted q-values < 0.05 were considered statistically significant.

### Single-sample gene set enrichment analysis (ssGSEA)

The background corrected and quantile normalized expression matrix was used as input for ssGSEA^59,60^ using a custom table of adaptive stress response (ASR) pathways previously described^29,31^. For each pathway, an estimated marginal mean was computed across AOIs using the pathway enrichment score. Patient was included as a random intercept and slide as a fixed effect (score ∼ Tissue_type + slide + (1 | Patient))^61^ in each tissue type using emmeans v2.0.3^62,63^. A pairwise contrast between tissue types was compared, and multiple testing correction (BH-FDR)^64^ applied within each contrast and across all ASR pathways.

### Cell Culture and Treatments

SUM149PT cells obtained from Asterand Inc. (Detroit, MI), MDA-IBC3 (PDX) cells cultured as described previously^65^ and TLAB-CC were collected as described in (Way et al., 2026)^31^ were cultured in Ham’s F12 medium (Mediatech Inc., Manassas, VA) supplemented with 5% FBS (Atlanta Biologicals, Lawrenceville, GA), 1% penicillin/streptomycin, 1% antibiotic/antimycotic, hydrocortisone (Invitrogen, Carlsbad, CA) and a human recombinant insulin in zinc solution (Gibco, Carlsbad, CA) at 37°C with 5% CO_2_ for no more than 2 months prior to use in this study. Cells were treated for 24 h in regular growth medium with Disulfiram (Sigma), CuSO_4_ (VWR), and DMSO as vehicle control. 10mM stock solutions of DSF (Allensworth et al., 2015) were made in dimethyl sulfoxide (DMSO) and subsequently diluted in media to concentrations as indicated. 10mM stock of CuSO4 was made in the media. Treatments as shown were carried out after culturing the cells overnight. Cell viability was determined by cell titer glow assay (Promega) where normalized % cell viability was calculated as luminescence with treatment/ luminescence with Vehicle control ×100. For caspase assay, the dye (Sartorious, Cal# 4440) was added to the culture media along with positive control (Staurosporine/Stauro;10µM), vehicle, or DSF [500nM] in presence of Cu [10µM]. Caspase activity was measured in cells following 22Hr of treatment as area under the curve (AUC) for normalized to untreated and compared with the control.

### Evaluation of DSF-Cu Treatment in the T-LAB Platform

To evaluate the effects of disulfiram (DSF; 250 nM) and copper (Cu; 10 µM) on tumor cell viability and invasion under lymphatic-like conditions, SUM149-GFP-Luc cells were cultured using the T-LAB platform. Briefly, cells were pretreated with DSF-Cu for 6 h before seeding. Transwell inserts were primed with serum-free medium containing DSF-Cu for 15 min, followed by the addition of 75 µL collagen (300 µg/mL) and incubation at room temperature for 1 h. Subsequently, 2.5 × 10⁵ pretreated live SUM149-GFP-Luc cells were seeded into the transwell inserts in 200 µL serum-free medium containing DSF-Cu. The T-LAB system was assembled with 6 mL serum-containing medium supplemented with DSF-Cu, and fluid circulation was maintained using a peristaltic pump (setting 1). The platform was maintained at 37°C under continuous flow and gentle agitation (Belly Dancer, setting 4) for 3 days. An additional 500 µL of serum-containing medium supplemented with DSF-Cu was added to the lower compartment on Days 1 and 2. On day 3, Cell viability of the cells present in cell suspension from transwell, collagen layer in the trans-well and in lymph-like fluidic part were assessed with trypan blue exclution assay using countess and the final cell viability was calculated as percentage of 2.5 × 10⁵ pretreated live SUM149-GFP-Luc cells.

## Supplementary Tables

**Supplementary Table 1.** The differentially expressed genes (DEGs, de_call = “up” or “dn”. Non-DEGs, de_call = “no”) identified in all contrasts for the GeoMx Spatial Profiling comparisons between adjacent normal (normal) tumor stroma (mid) edge and lymphovascular (LVI) compartments. Related to Figure 3.

**Supplementary Table 2.** Gene Set Enrichment Analysis (GSEA) using the Hallmarks of Cancer gene sets (www.gsea-msigdb.org, H) for all contrasts of the GeoMx Spatial Profiling comparisons. Related to Figure 4.

**Supplementary Table 3.** Gene Set Enrichment Analysis (GSEA) using the Gene Ontology: Biological Processes gene sets (www.gsea-msigdb.org, C5, subcollection GO) for all contrasts of the GeoMx Spatial Profiling comparisons. Related to Figure 5 and 6.

**Supplementary Table 4.** Gene Set Enrichment Analysis (GSEA) using the Gene Ontology: Molecular Function gene sets (www.gsea-msigdb.org, C5, subcollection GO) for all contrasts of the GeoMx Spatial Profiling comparisons. Related to Figure 5 and 6.

**Supplementary Table 5.** Gene Set Enrichment Analysis (GSEA) using the Gene Ontology: Cellular Components gene sets (www.gsea-msigdb.org, C5, subcollection GO) for all contrasts of the GeoMx Spatial Profiling comparisons. Related to Figure 5 and 6.

**Supplementary Table 6.** The shared pathways (GO:BP, GO:CC, GO:MF separate tabs) enriched in SUM149 and TLAB-CC comparisons and in the corresponding tissue compartments (stroma corresponding to SUM149, LVI corresponding to TLAB-CC) of the GeoMx Spatial Profiling dataset. The column “same_direction” indicates the pathway was enriched in the same direction in both comparisons. Related to Figure 6.

